# 3-Minute Hematoxylin and Oil Red O (H-ORO) Staining Protocol for Frozen Sections of Zebrafish

**DOI:** 10.64898/2026.04.03.716422

**Authors:** Choy Kim, Seong-Kyu Choe, Seok-Hyung Kim

## Abstract

Optimized histological techniques are crucial for visualizing cellular morphology across zebrafish tissues. Here, we report a rapid and reliable hematoxylin and Oil Red O (H-ORO) staining protocol for frozen sections that can be completed in less than three minutes. Mayer’s hematoxylin is used for nuclear staining, followed by Oil Red O (ORO) to visualize lipid-rich structures such as the endomysium surrounding myofibers, white matter of the brain, and myelin layers of major axonal tracts. Importantly, our optimized H-ORO protocol preserves tissue integrity and minimizes artifacts such as myofiber shrinkage commonly observed with ethanol-based hematoxylin and eosin (H&E) staining in both frozen and paraffin sections.

## Introduction

Hematoxylin and eosin (H&E) staining remains the standard method for histological analysis of tissue morphology (1, 2). In zebrafish research, frozen sections embedded in agarose are more commonly used than paraffin or plastic sections. However, ethanol-based dehydration steps and xylene mounting required for traditional H&E staining often disrupt cellular architecture in frozen tissues. In particular, dehydration-induced shrinkage of myofibers is a frequent artifact that can lead to misinterpretation of pathological changes. To address these limitations, we optimized a rapid, water-based histological staining method for frozen zebrafish sections that preserves morphological detail and can be completed in less than three minutes.

### Materials and Methods Zebrafish husbandry

All zebrafish were maintained under standard laboratory conditions following the guidelines of the Committee for Ethics in Animal Experiments at Wonkwang University. Experimental procedures were approved by the Institutional Animal Care and Use Committee (Wonkwang University IACUC #WKU23-77).

### Reagents

Mayer’s modified hematoxylin solution (Abcam, AB220365) was used for nuclear staining, and 0.5% Oil Red O (ORO) in isopropanol (Sigma, 01391) for lipid staining.

### Preparation of frozen sections

Larval or adult zebrafish were fixed overnight in 4% paraformaldehyde (PFA) at 4°C. Adult zebrafish were euthanized in ice water and then cut into three segments to minimize internal tissue damage by pancreatic digestive enzymes. Fixed samples were embedded in 1.2% agarose containing 5% sucrose (w/v), then cryoprotected in 30% sucrose overnight. Sucrose buffer was removed by paper towel and blocks were put on the plastic mold. The plastic mold was placed on a frozen stainless-steel container which was inserted in a Styrofoam box with liquid nitrogen. Cryosections (8 μm thickness) were cut at -30°C.

### H-ORO staining procedure

1. Thaw the slide for 30 s and rinse with distilled water (DW).
2. Immerse the slide in Mayer’s hematoxylin (in a slide mailer) and stain for 1 min.
3. Rinse gently with DW using a squeeze bottle.
4. Submerge the slide in bluing buffer (0.1% sodium bicarbonate) for 20 s.
5. Rinse again with DW.
6. Remove excess water and place the slide on a paper towel.
7. Prepare fresh ORO working solution just before use: mix 200 μL DW with 300 μL of ORO stock (0.5% ORO in 60% isopropanol) in a 1.5 mL microtube and mix vigorously to prevent precipitation.
8. Apply 150–200 μL of ORO working solution to the slide and stain for 20 s.
9. Rinse with DW.
10. Mount using 75% glycerol in PBS.
11. Proceed to imaging.

### Imaging

Images were acquired using an upright microscope (Zeiss, Primostar 1) at the Core Facility for Supporting Analysis & Imaging of Biomedical Materials at Wonkwang University supported by National Research Facilities and Equipment Center.

## Results

The optimized H-ORO protocol enables complete histological staining of agarose-embedded frozen zebrafish sections within three minutes (Fig. 1a). The method yields high-contrast nuclear and lipid-enriched structure staining with minimal tissue cracking or shrinkage in 7 days post-fertilization (dpf) larvae (Fig. 1b). Following feeding, ORO staining clearly delineated lipid-rich blood vessel structures in larval brains (Fig. 1c).

**Figure 1.**
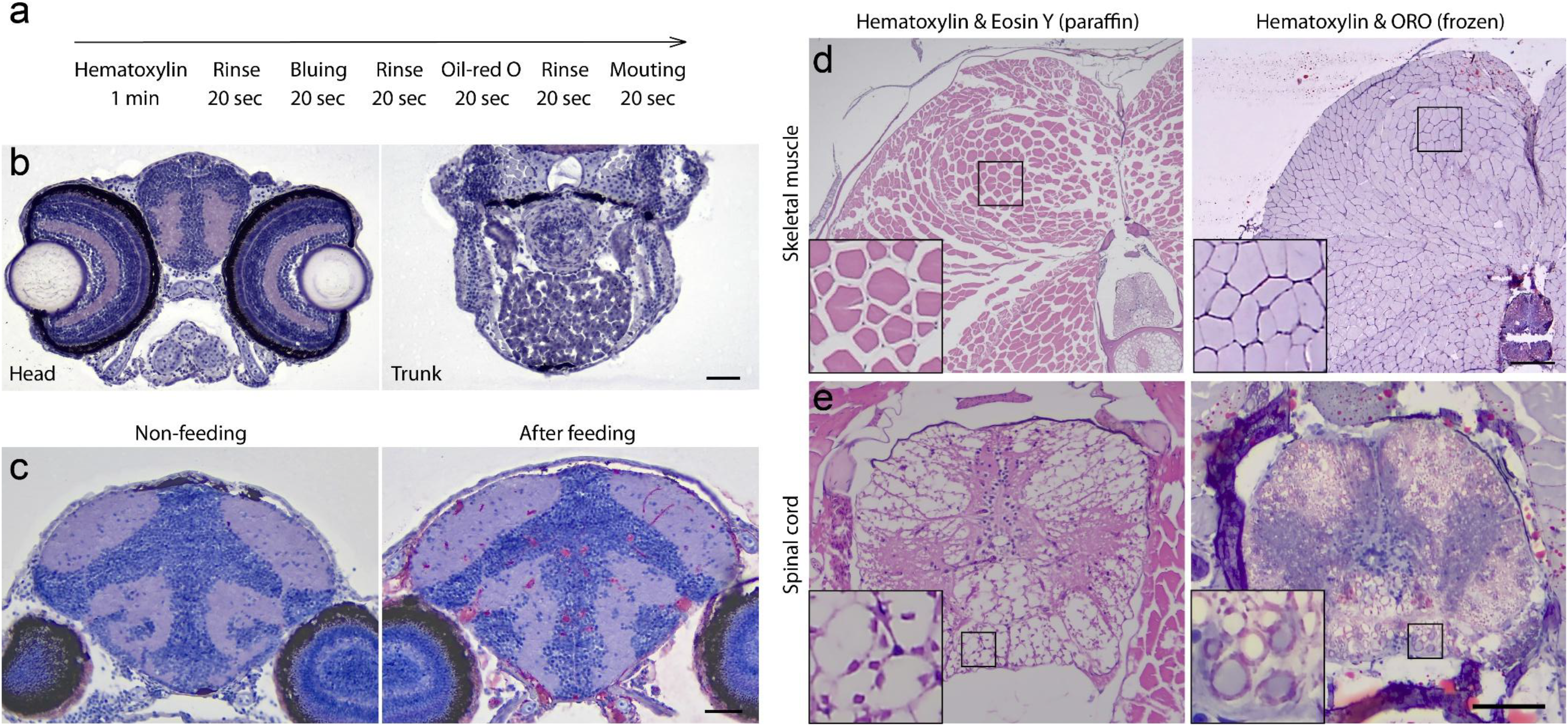
H-ORO staining in agarose-embedded frozen sections of larval and adult zebrafish. (a) Workflow schematic of the H-ORO staining protocol. (b, c) Representative images H-ORO staining from head and trunk sections of 7 dpf larvae. Scale bar = 50 µm. (d) Comparison of skeletal muscle morphology between H&E (paraffin) and H-ORO (frozen) sections. Scale bar = 250 µm. (e) Spinal cord staining in paraffin (H&E) vs. frozen (H-ORO) sections. Scale bar = 50 µm.

Compared to conventional ethanol-based H&E staining on paraffin sections, which commonly causes myofiber shrinkage and lipid loss, H-ORO preserved muscular morphology and myelin-rich white matter in the spinal cord (Fig. 1d, e). This demonstrates superior preservation of lipid-containing structures and overall morphological integrity.

## Authors’ Contributions

CK performed experiments, SKC edited the manuscript and funding acquisition. SHK conceived the study, performed experiments, wrote the manuscript, and acquired funding.

